# Live cell monitoring for factors affecting genome variation

**DOI:** 10.1101/508150

**Authors:** Yuntao Xia, Kuangzheng Zhu, Jerome Irianto, Jason C. Andrechak, Lawrence J. Dooling, Charlotte R. Pfeifer, Dennis E. Discher

## Abstract

Cancer cells and pluripotent stem cells frequently exhibit gains or losses of entire chromosomes and chromosome segments, and the typical terminal analyses of genomes suggest this aneuploidy is ongoing and particularly variable in solid tumors. Here, we quantify aneuploidy-inducing perturbations by live cell fluorescence monitoring for changes in chromosome-5 in a lung cancer line and in normal diploid iPS cells. Inhibition of the spindle assembly checkpoint (SAC) and knockdown of DNA repair factors cause chromosome mis-segregation to increase several-fold above a low baseline level, and both perturbations also generate several-fold more rare fluorescent-null cells. Loss of chromosome-5 is confirmed by single cell karyotyping, SNP arrays on stable isolated clones, and downregulated expression of genes on chromosome-5 in single cell transcriptomics. The iPS cells also show loss of fluorescence in infrequent cells after SAC inhibition and upon growth as teratomas in mice. Viability, selection, and altered expression can thus be tracked to reveal molecular mechanisms in aneuploidy.

## Introduction

Copy number changes are a major form of genomic variation in cancer (Hastings et al., 2009; Zhang et al., 2009) and are often more common than somatic mutations. A recent study of medulloblastoma, for example, shows the most frequently detected mutation (in *MYC*) occurs in only 17% of patients whereas every one of more than a dozen copy number changes occur in more than 30% of patients (Northcott et al., 2017). In clinical trials by Yamanaka and colleagues on two aged patients (>65 yrs), induced pluripotent stem (iPS) cells generated from one of the patients exhibited copy number variations that prevented their use for fear of causing cancer (Mandai et al., 2017). Mechanisms for copy number variation remain unclear, and reported changes are likely to reflect selection and affect gene expression (i.e. gene dosage).

Current methods to assess copy number variations require DNA extractions for sequencing and arrays or else viable mitotic cells for metaphase spreads, but such methods are terminal experiments for cells. Rare changes (~1% of cells) are also difficult to be confident in, and whether a given change yields a viable cell can at best be inferred. A live cell reporter for copy number changes in rare cells might address such issues and be useful to decipher mechanisms. A recently reported live reporter in yeast has used changes in intensity of a chromosome-integrated, GFP-tagged gene to hint at copy number changes, but validations and imaging were limited (Lauer et al., 2018). In various human cancers, many chromosomes are lost or gained, and loss of Chr-5 is commonly seen in gastric, esophageal, lung, ovarian and breast cancer (Johannsdottir et al., 2006; Mendes-da-Silva et al., 2000; Michelassi et al., 1989; Ogasawara et al., 1996; Tavassoli et al., 1996). Here, we describe a chromosome-5 (chr-5) reporter in live human-derived cells with detailed copy number characterizations and perturbations, followed by corresponding gene expression assessments and sample studies both *in vitro* and *in vivo*.

## Results

### Loss of chromosome 5 reporter in live cells

We hypothesized that fusing a GFP variant protein to a housekeeping gene in one allele of Chr-5 will allow us to look for loss of the fluorescent signal and relate to loss of all or part of Chr-5 (Fig. 1A). The lung cancer line (A549) is hypotriploid, and one of the three copies of chr-5 is gene-edited with RFP fused to the N-terminus of a housekeeping protein while protein from the other two chr-5s is unaltered (Fig. S1A). Modified cells all express RFP signal after cell sorting. After weeks in standard culture, however, some RFP-neg cells were evident as viable colonies with no obvious growth defects (Fig. 1B). RFP-neg cells were clonally expanded for genomic characterization in order to test our hypothesis.

**Figure 1.**
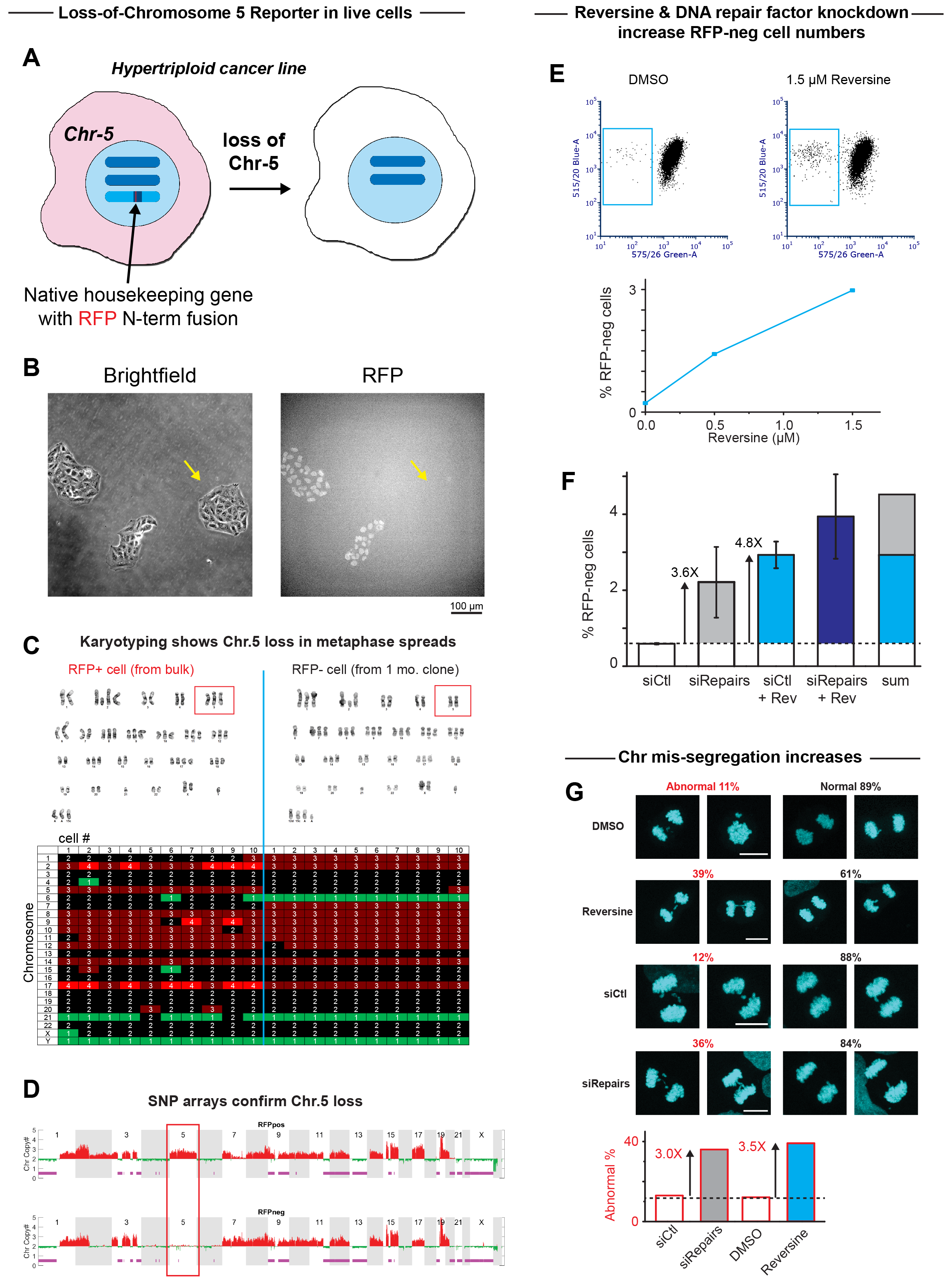
(A) A housekeeping gene on one of the Chr-5s is fused with a RFP sequence at the N terminus, so that cells (A549) will be RFP-pos when the protein is expressed. RFP signal will disappear when the modified chr-5 is lost. (B) Viable colonies are seen among RFP-pos and RFP-neg cells. (C) Karyotyping shows chr-5 loss in RFP-neg cells based on metaphase spreads. Heterogeneity is observed when comparing 10 metaphase spreads of bulk RFP-pos cells. Clonal RFP-neg cells after one-month culture exhibit some level of heterogeneity such as a loss in chr-12 (cell#1) and a reverter in chr-5 (cell#10). (D) SNP arrays confirm chr-5 loss in RFP-neg cells. (E) Reversine treatment leads to increased RFP-neg cell numbers in a dose-dependent manner. (F) Knockdown of DNA repair factors increases RFP-neg cell numbers. Combining reversine and DNA repair knockdown has an additive effect suggesting two pathways may function orthogonally to maintain genomic stability. (G) Aneuploidy is seen at ~10% among A549 during mitosis. However, reversine treatment and knockdown of DNA repair factors cause ~3 times more occurrence of aneuploidy, consistent with the fold change in RFP-neg cells.

Karyotyping of ten metaphase spreads showed 90% of RFP-neg cells with only two copies of chr-5 whereas RFP-positive cells always showed three copies (Fig. 1C). Karyotype variation in the bulk population of RFP-pos otherwise reflects the genomic instability of these cancer cells (Fig. 1C), and even in clonal RFP-neg cells, the one month of culture needed for clonal expansion reveals a loss of chr-12 in cell #1 and a revertant gain of chr-5 in cell #10 (Fig. 1C). To confirm the loss of RFP, PCR used one primer in the RFP region and a second primer in the native fused gene: no band is detected in RFP-neg cells compared to RFP-pos, and the sensitivity is ~1:1000 or better (Fig. S1B).

SNP (Single Nucleotide Polymorphism) array analysis for copy number changes (Irianto et al., 2017) again reveal chr-5 loss in RFP-neg clones (Fig. 1D). A few differences between karyotyping and SNP arrays include only one chr-6 in metaphase spreads of RFP-neg cells (Fig. 1C,D), but detailed analyses indicate a translocation of chr-6 to chr-1. Unidentified chromosomes in karyotyping will also be detected in SNP arrays such as the gain of chromosome segments on chr-3 (Fig. S1C). Regardless, after correction and subtraction of RFP-pos from RFP-neg, the two methods agree with each other (Fig. S1C-bottom) and consistently show loss of RFP signal in cells correlates with a loss of chr-5 (Fig. 1C, D, S1C-bottom).

### Reversine & DNA repair factor knockdown increase RFP-neg cell numbers

In normal cells, mis-segregation of chromosomes is prevented by the spindle assembly checkpoint (SAC), which inhibits mitotic exit until all chromosomes have achieved bipolar attachment to spindle microtubules (Barbosa et al., 2011; Lara-Gonzalez et al., 2012). Reversine perturbs SAC and drives appearance of RFP-neg cells in a dose-dependent manner easily quantified by flow cytometry (Fig. 1E), which suggests a loss of chr-5. Depletion of DNA repair factors upon nuclear envelope rupture causes excess DNA damage and might also produce copy number changes (Irianto et al., 2017; Xia et al., 2018). Knockdown of multiple DNA repair factors (i.e. KU80, BRCA1, and BRCA2) indeed drives increased genomic heterogeneity in the samples (Fig. S2) and ~3-fold more cells become RFP-neg (Fig. 1F). Combination of this knockdown with reversine suggests an additive effect (Fig. 1F), consistent with orthogonal functions of SAC and DNA repair in genomic stability.

Dividing cancer cells exhibit ~10% abnormalities in a standard assay for lagging chromosomes (Fig. 1G), which may explain the appearance of ~0.5% RFP-neg cells in long-term cultures of pure RFP-pos cells (Fig. 1E,F). However, such aneuploidy increases to nearly ~40% (~3 fold) after reversine treatment or knockdown of DNA repair factors (Fig. 1G). The fold increase in mitotic errors is consistent with the fold-increase in RFP-neg cells under the same perturbation (Fig. 1F).

### Gene expression correlates with chromosome numbers

Cell-to-cell variations in epigenetic states generally add complexity to gene expression profiles, but copy number variation seem detectable in transcript profiles per a ‘gene dosage’ effect (Raznahan et al., 2018; Gao et al., 2016). To minimize complexities of cell-to-cell variation, single cell RNA-seq was applied to four RFP-neg clones and four RFP-pos clones as two mixtures. Chromosome copy numbers of eight clones were also assessed by SNP array and again show loss of one copy of three chr-5’s in RFP-neg cells (Fig. S3). Single cell RNA-seq showed 7385 genes account for >95% of total reads, which were normalized by Transcripts Per Kilobase Million (TPM) (Hwang et al., 2018; Vallejos et al., 2017). Comparing RFP-neg and RFP-pos cells reveals a Chr-5 specific, chromosome-wide decrease in gene expression (by ~1/3^rd^) at both single cell and bulk levels (Fig. 2A).

**Figure 2.**
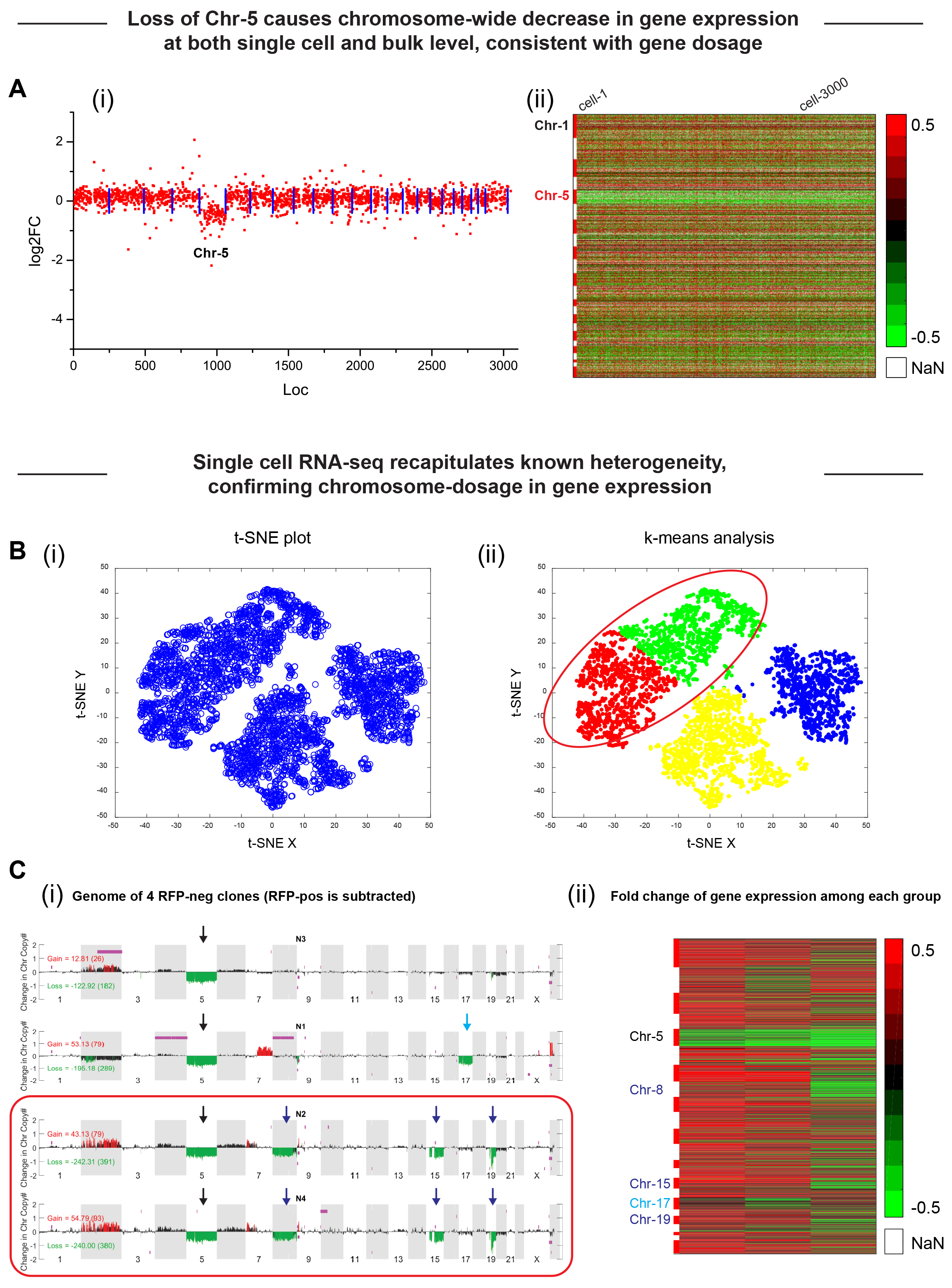
(A) After calculating TPM and the ratio of RFP-neg/RFP-pos of 7385 genes, RFP-neg cells show chromosome-wide decrease in gene expression at both the bulk and single cell levels, consistent with the gene dosage hypothesis. (B) Combing t-SNE and k-means analysis, single cell RNA-seq recapitulates known heterogeneity in the original samples. (C) The gene expression of each subpopulation matches well with corresponding genotype. The box labels two clones that have identical genome, which explains the mingled population in B-ii.

Single cell RNA-seq has been applied to distinguish different cell types in tissues as these cells typically have distinct gene expression profiles, and the results can be validated by means such as marker-specific immunostaining (Przepiorski et al., 2018). RFP is the principal marker here, but heterogeneity due to the mixture of four RFP-neg clones can also be identified using t-SNE and k-means analysis (Fig. 2B) (Grün et al., 2015). A large intermingled population in t-SNE and k-mean plots is likely due to two RFP-neg clones having an identical genome (Fig. 2B-ii&C-i). However, loss of chr-17 in a separate population and loss of chr-8, 15 and 19 in the another population correlate well with each genotype as well (Fig. 2Ci-ii). The “chr-5 reporter system” thus suggests that single cell RNA-seq can clarify gene dosage and separate cancer cell sub-populations with minor difference in genotypes.

### Phenotypic differences can also associate with genotypic differences

Although the four RFP-pos clones exhibit similar genotypes, clone-1 uniquely gains a chromosome 7 q-arm (Fig. 3A). The copy number changes in chromosome 7 have been seen in many diseases such as 7q11.23 duplication syndrome and Williams-Beuren syndrome, which causes delayed cell growth and neurological problems (Berg et al., 2007; Sanders et al., 2011). Cells in clone-1 exhibit slightly delayed proliferation and less spindle-polraized morphology (Fig. 3B,C). As cell migration requires polarization (Ridley et al., 2003), live imaging was done on sparse culture. Clone-1 proves to be less migratory than any other clone, with no directional bias obvious in any clone (Fig. 3D). Although further bioinformatic analysis is required to resolve mechanism(s) linking genotype and phenotype, we had shown in a similar study that an extra copy of Chr-8 increases GATA4, which promotes microtubule assembly, spindle shapes, and faster migration in 3D (Irianto et al., 2017).

**Figure 3.**
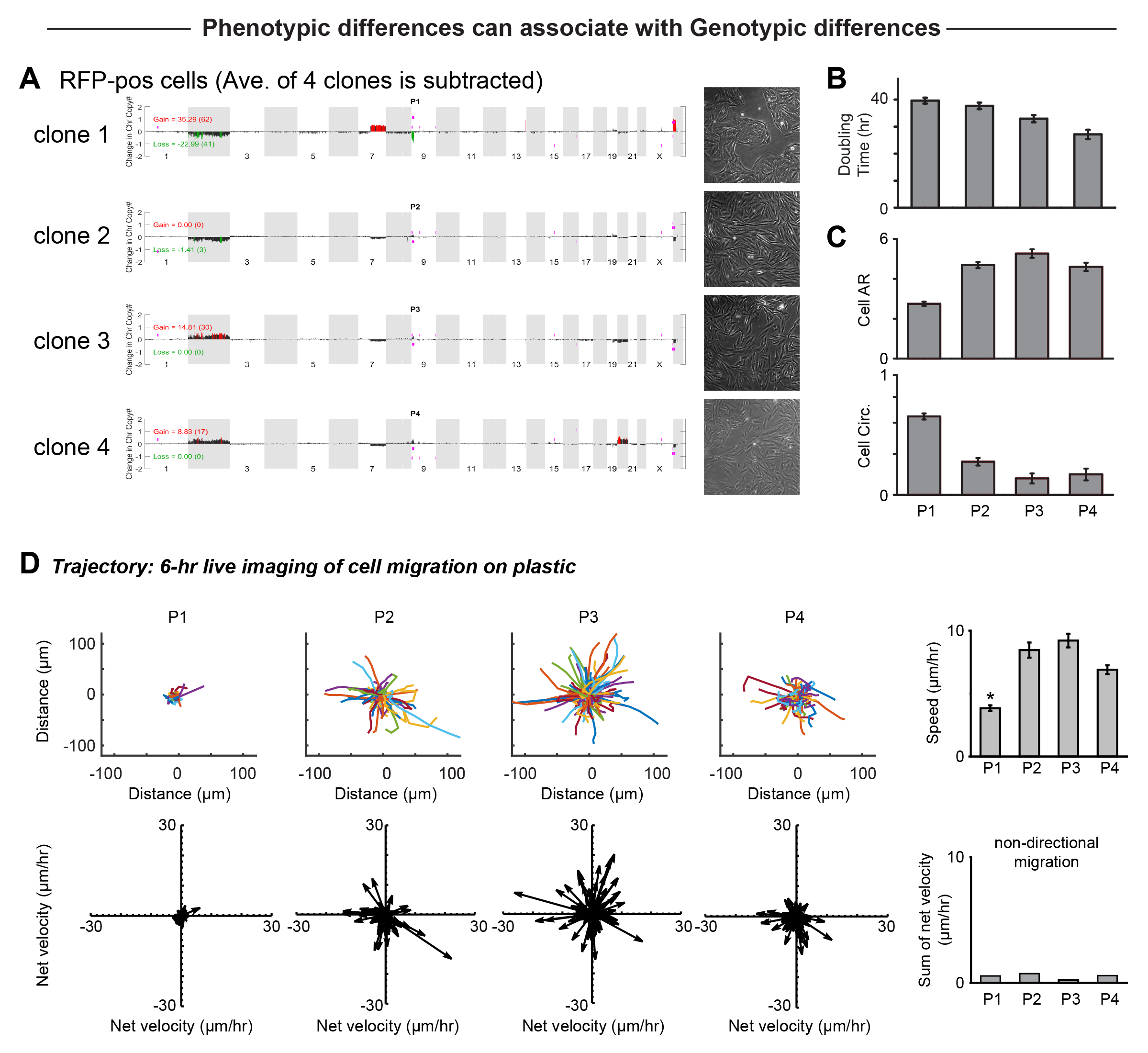
(A-C) Comparison between 4 RFP-pos clones indicates clone-1 has a gain on chr-7 q-arm. Clone-1 cells are more rounded, proliferate slightly slower, and grow in clusters. (D) Live imaging of the 4 clones shows clone-1 is less migratory. No directional preference in migration is observed in any clone.

### Normal diploid iPS cells may lose chromosomes *in vitro* and *in vivo*

Cancer cells typically possess intrinsic genomic instability and tend to gain or lose chromosomes (Negrini et al., 2010) – such as chr-5 in culture (Fig. 1A,B). In comparison, iPS cells derived from a young adult human with similar gene editing of chr-5 are stable, showing 100% GFP-pos after weeks of culture. SNP array analysis of bulk cultures also show no copy number variation. Synchronizing cell cycle of iPS cells by nocodazole causes severe cell death and the remaining survivors do not divide even after release. However, iPS cells survived reversine treatment for a few passages, and some GFP-pos iPS cells became GFP-neg based on both flow cytometry and imaging (Fig. 4A).

**Figure 4.**
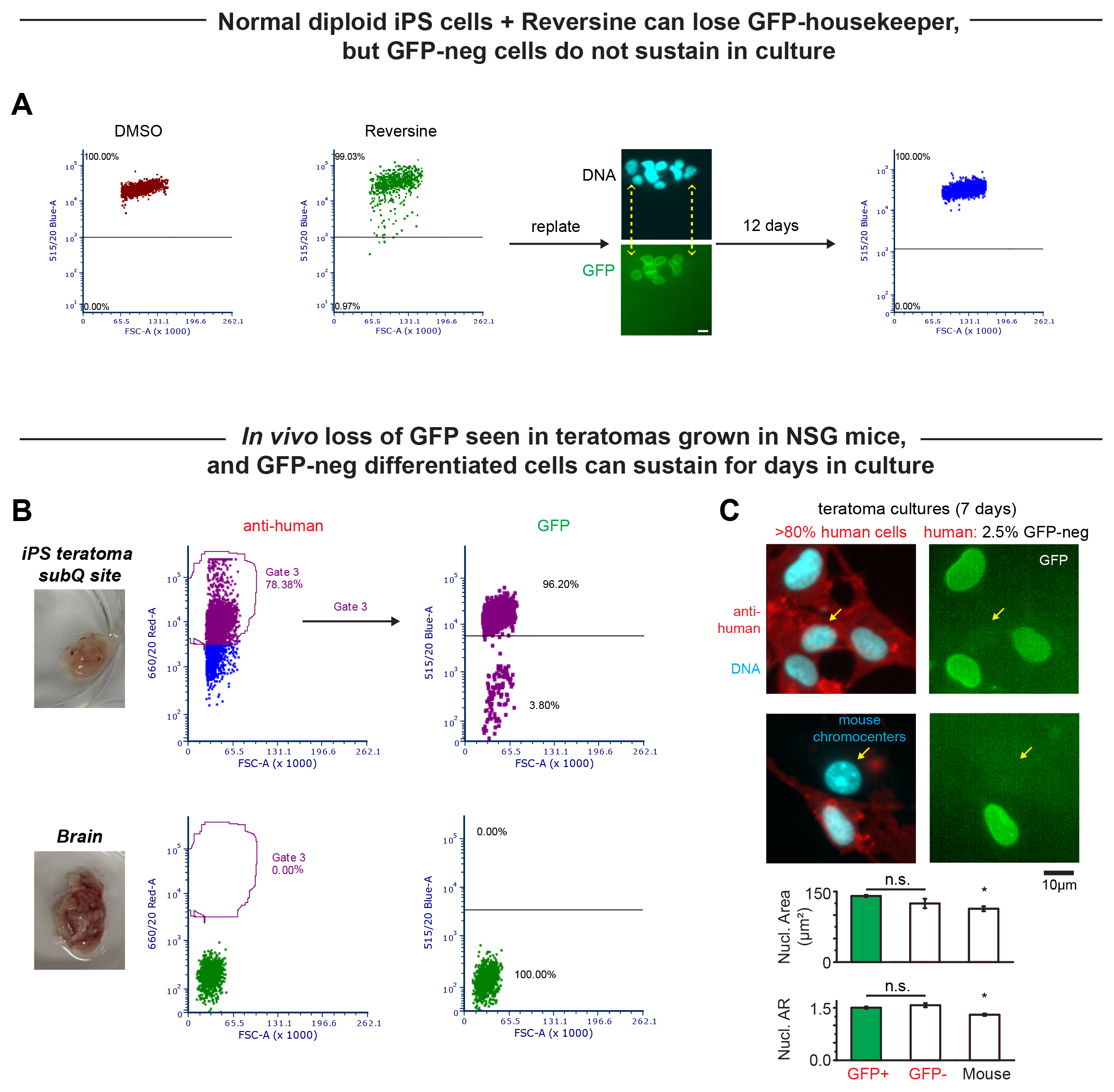
(A) Normal diploid iPS cells with similar gene editing on chr-5 shows 100% GFP-pos. However, loss of GFP signal is seen after reversine treatment, confirmed by both flow cytometry and microscopic images. These GFP-neg iPS cells suffer from survivability after days of culture. Scale bar: 10 μm. (B) iPS cells grow into teratomas in subcutaneous sites in mice. GFP-neg cells *in vivo* are confirmed by flow cytometry and immunofluorescence.

Our GFP-pos iPS cells grow robustly in clusters and double every 12 hrs in standard culture, but GFP-neg iPS cells do not persist (Fig. 4A). This is probably because 1) normal cells intrinsically lose survivability after chromosome loss (Teng and Hardwick, 2015), and 2) a standard culture environment on Matrigel-coated plastic is not suitable to support genomically-aberrant cells. In particular, aged mouse-iPS cells with genomic instability can be diverted into apoptosis by improving the DNA damage response with ‘simple’ addition of a cell-permeable antioxidant during reprogramming (Skamagki et al., 2017). This exciting, recent finding did not address, however, whether abnormal genomes in the source cells (eg. fibroblasts) are the origins of the genomic changes in the iPS cells – as often proposed.

Teratomas in an NSG mouse were made by subcutaneous flank injections of the iPS cells following other studies (Gutierrez - Aranda et al., 2010; Nelakanti et al., 2015). After 4 wks, cells dissociated from teratoma and other tissues such as brain were analyzed by flow cytometry or else cultured on Matrigel-coated plastic. Mouse and human cells can be clearly distinguished by anti-human antibody based on control studies with mouse melanoma-derived B16F10 cells (Fig. S4A). Using the same staining and flow cytometry settings, GFP-neg iPS cells were detected by both flow cytometry and imaging (Fig. 4B,C). Cells seem to have differentiated already as abundant lamin-A is seen and they no longer grow in clusters (Fig. S4B). All isolated GFP-neg cells from the teratoma again died by 2 wks in culture, even though most of the isolated GFP-pos iPS cells remain viable in culture. Nonetheless, the appearance of GFP-neg iPS cells at least suggest some stresses *in vivo* are sufficient to cause aneuploidy among normal iPS cells – even though loss of all of part of or the entire chr-5 must be assessed by methods above.

## Conclusion

The chromosome reporter in live cells demonstrated here has the potential for the study of factors that cause copy number changes in both cancer cells and normal cells. Although we only show chr-5 loss in our characterization, chr-5 gain is also detectable in theory. Ultimately, if one can label all chromosomes with different colors, then all chromosomes can be monitored simultaneously. Although there is a correlation between copy number changes and gene expression, the change is usually ~30-50% when a gain or loss of the chromosome is seen. Many cancers cells indeed express markers that are many times more than that in normal cells, which may not be a simple effect caused by chromosome changes, but more complicated secondary responses may be involved to cause the issues and could be illuminated by the approaches here.

## Material and Methods

### Cell culture

A549 cells were cultured in F12 media (A549), supplemented with 10% FBS and 1% penicillin/streptomycin under 37°C and 5% CO_2_ as recommended by ATCC. When drug treatment is required, 1.5μM reversine was used for 24 hrs, followed by releasing for 48 hrs. If imaging on mitotic cells is needed, after DMSO or reversine treatment, the media was replaced with 1.5μg/ml Nocodazole for 18 hrs and released in pure media for 1 hr.

All iPS cells were cultured in mTeSR1 medium, supplemented with 5X growth factors, incubated at 37°C and 5% CO_2_. Plastic well plates were coated with Matrigel in DMEM media for 1 hr before cells were plated.

### Immunostaining

Cells were fixed in 4% formaldehyde (Sigma) for 15 minutes, followed by permeabilization by 0.5% Triton-X (Sigma) for 15 minutes, and blocked with 5% BSA (Sigma) for 30 minutes and overnight incubation in primary antibodies (anti-human). The cells were then incubated in secondary antibodies (1:500 ThermoFisher) for 1.5 hours, and their nuclei were stained with 8μΜ Hoechst 33342 (ThermoFisher) for 15 minutes. When mounting is involved, Prolong Gold antifade reagent was used (Invitrogen, Life Technologies).

### Imaging

Epifluorescence imaging was performed using an Olympus IX71 with a digital camera (Photometrics) and a 40X/0.6 NA objective. Confocal imaging was performed on a Leica TCS SP8 system with a 63X/1.4 NA oil-immersion. Live imaging was performed on an EVOS FL Auto Imaging System with 10× or 20× object in normal culture conditions (37°C and 5% CO_2_; complete culture medium as specified above).

### Transfection/Knockdown

All siRNAs used in this study were purchased from Dharmacon (ON-TARGETplus SMARTpool siBRCA1, L-003461-00; siBRCA2, L-003462-00; siKu80, L-010491-00). A549 cells were passaged 24 hours prior to transfection. A complex of siRNA pool (25 nM each) and 1 μg/mL Lipofectamine 2000 was prepared according to the manufacturer’s instructions and incubated for 3 days (for siRNAs) in corresponding media with 10% FBS. Knockdown efficiency was determined by Immunoblot following standard methods (Xia et al., 2018).

### Flow cytometry / Sorting

For flow cytometry, A549 cells were dissociated using 0.05% trypsin for 5 min, washed in PBS, and re-suspended in 5% FBS in PBS. Samples were run on a BD LSRII Flow Cytometer.

For sorting, cells were prepared in the same way as flow cytometry, except that cells were kept sterile. FACS Aria (BD Biosciences) was used to sort. Sorted cells were collected in 15ml centrifuge tubes, with 10% FBS and 1% P/S in corresponding media (complete media as described in cell culture section). Sorted cells were then centrifuged with 3000 rpm for 5 min and cultured back in T25 or T75 flasks.

If immunostaining is need for flow cytometry, cells were washed with PBS after trypsinization, resuspended in 5% BSA in PBS and put onto a rotator for 15min. Cells were then centrifuged, supernatant discarded; the same process was applied with 1:500 FC receptor in 5% BSA. Next, cells were resuspended in 1:500 primary antibody (anti-human), left on rotator for 20 min, centrifuged, supernatant discarded, and washed once with 1%FBS in PBS. The same process was repeated for secondary antibody, also at 1:500 dilution. Finally, cells were resuspended in 5% FBS with PBS, and run on flow cytometer as described before.

### Karyotyping

Cells used for karyotyping were plated in T25 flasks, cultured for 2-3 days to reach half confluency. The media was then discarded and replaced with new media to fill the entire flask with a closed lid, wrapped with parafilm. The samples were then sent to a company to conduct karyotyping.

### Genome (SNP array) analysis

DNA was isolated with the Blood & Cell Culture DNA Mini Kit (Qiagen) per manufacturer’s instruction. The same DNA samples were sent to The Center for Applied Genomics Core in The Children’s Hospital of Philadelphia, PA, for Single Nucleotide Polymorphism (SNP) array HumanOmniExpress-24 BeadChip Kit (Illumina). For this array, >700,000 probes have an average inter-probe distance of ~4kb along the entire genome. For each sample, the Genomics Core provided the data in the form of GenomeStudio files (Illumina). Chromosome copy number and LOH regions were analyzed in GenomeStudio by using cnvPartition plug-in (Illumina). Regions with one chromosome copy number are not associated with LOH by the Illumina’s algorithm. Hence, regions with one chromosome copy number as given by the GenomeStudio are added to the LOH region lists. SNP array experiments also provide genotype data, which was used to give Single Nucleotide Variation (SNV) data. In order to increase the confidence of LOH data given by the GenomeStudio, the changes in LOH of each chromosome from each sample were cross referenced to their corresponding SNV data. After extracting data from GenomeStudio, all data analysis was done on Matlab.

### PCR

DNA was extracted as described in “SNP array section”. The isolated DNA was then mixed with materials from KAPA HiFi PCR Kit to start PCR: Each reaction contains 5μL 5X HiFi Fidelity Buffer, 0.75μL 10mM KAPA dNTP Mix, 0.5μL 1U/μL KAPA HiFi DNA Polymerase, 0.75μL of 10μM forward and reverse primers, respectively, and 1ng of extracted DNA template. PCR grade water was then filled up to 25μL. All materials suggested by the kit were placed on ice prior to mixing. The reaction mix was placed on the thermocycler with the following temperature cycling protocol: Initial denaturation at 95 °C for 3 min; 35 cycles of denaturation at 98 °C for 20 sec, annealing at 65 °C for 15 sec, extension at 72 °C for 15 sec; final extension at 72 °C for 1 min. Afterwards, PCR products were loaded on 1.2% agarose E-gel with Ethidium Bromide and run for 30 min. Images were taken by placing the gel under GENE Flashing UV source.

### Single cell RNA-seq

A549 cells from four positive clones and four negative clones were plated in 96 well plate at 150 K cells/well, respectively, and cultured for 2 days. The positive sample is a mixture of four positive clones, and the negative sample is a mixture of four negative clones. Two samples were submitted to Center for Applied Genomics (1014, Abramson Research Center, University of Pennsylvania) for single cell RNA sequencing. Analysis was done in either R or Matlab.

### Teratoma *in vivo*

For each injection, iPS cells were suspended in 300 μL ice-cold PBS and 30% Matrigel (BD) and injected subcutaneously into the flank of non-obese diabetic/severe combined immunodeficient (NOD/SCID) mice with null expression of interleukin-2 receptor gamma chain (NSG mice) (Alvey et al., 2017). Mice were obtained from the University of Pennsylvania Stem Cell and Xenograft Core. All animal experiments were planned and performed according to IACUC protocols. The teratoma were grown for 4 weeks and cells from teratoma and other tissues were then isolated for culture and flow cytometry.

### Statistics

All statistical analyses were performed using Excel (2013; Microsoft) or MATLAB. Unless otherwise noted, statistical comparisons were made by unpaired two-tailed Student’s t tests and were considered significant if p < 0.05. Unless mentioned, all plots show mean ± SEM. n indicates the number of samples, cells, or wells quantified in each experiment.

## Supplemental figure legends

**Figure S1.**
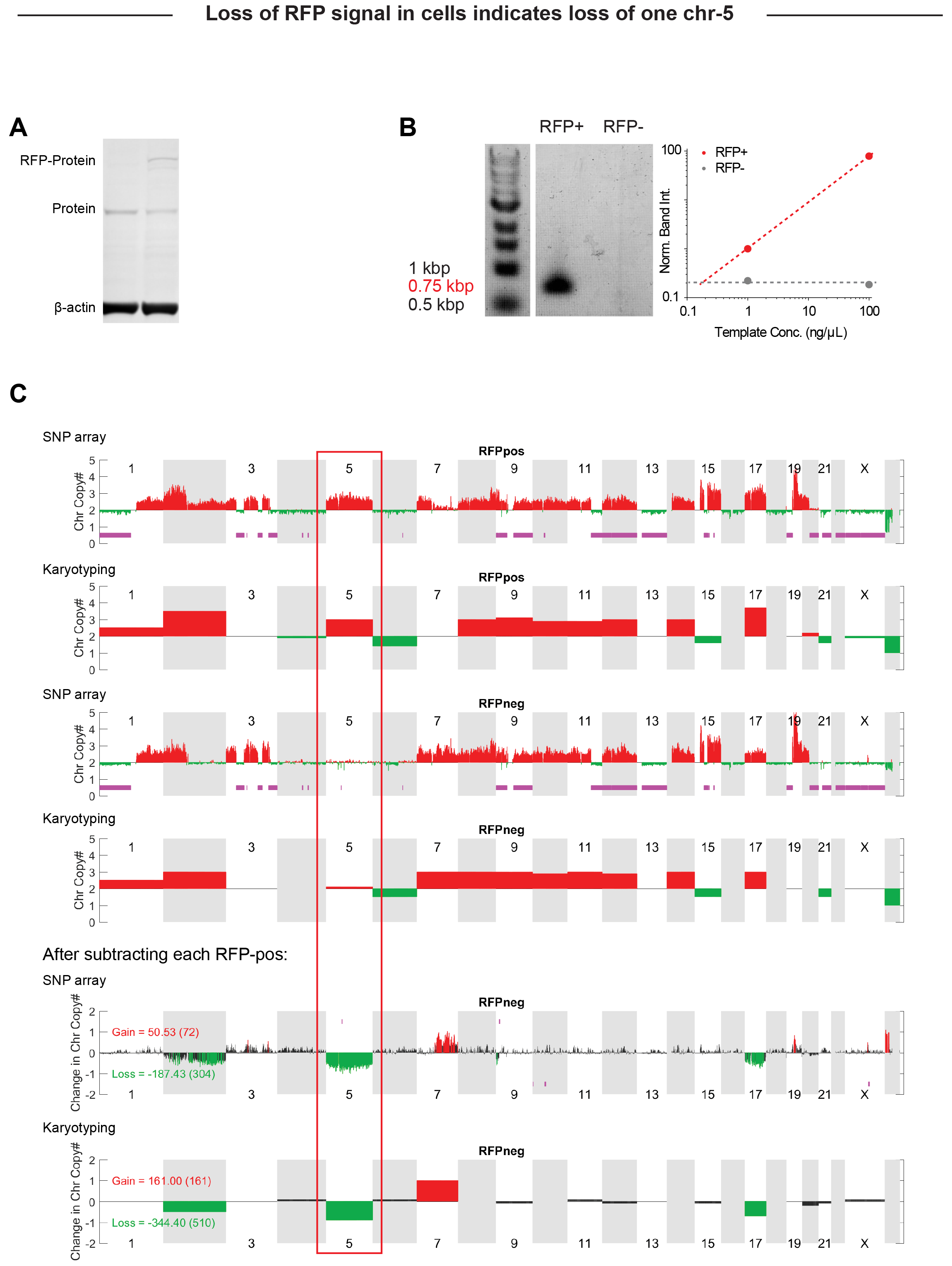
(A) Western blot of modified and unmodified A549 cells: As only one of the chr-5s is modified, endogenous protein without RFP can also be expressed. (B) PCR with one primer at RFP region and the other one in the gene region shows no band on RFP-neg samples. (C) SNP array and karyotyping consistently indicate loss of one chr-5 in RFP-neg cells.

**Figure S2.**
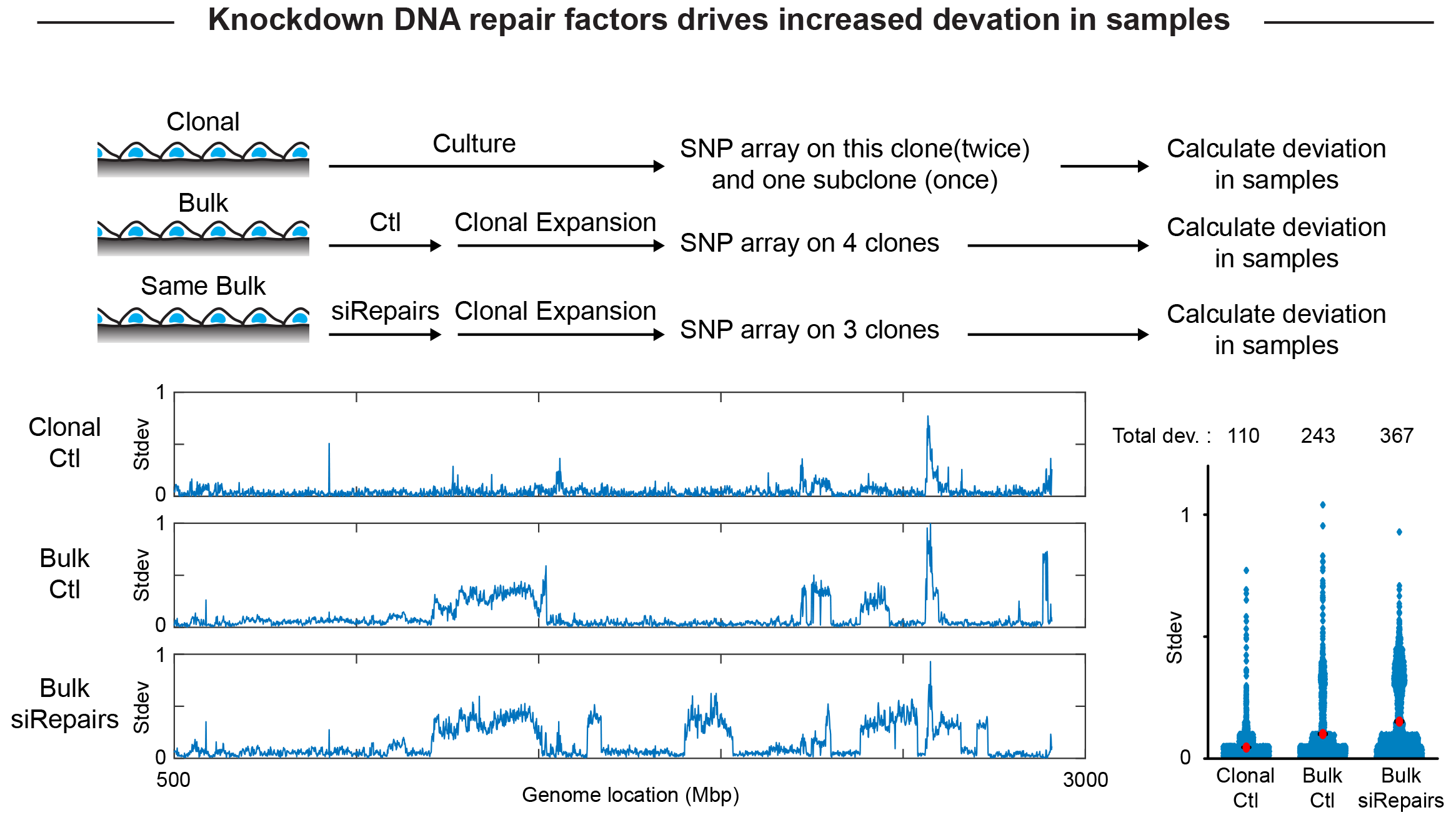
Schematic illustration of experimental setups. Knockdown of DNA repair factors drives increased deviation in SNP array reads between samples, suggesting genomic heterogeneity.

**Figure S3.**
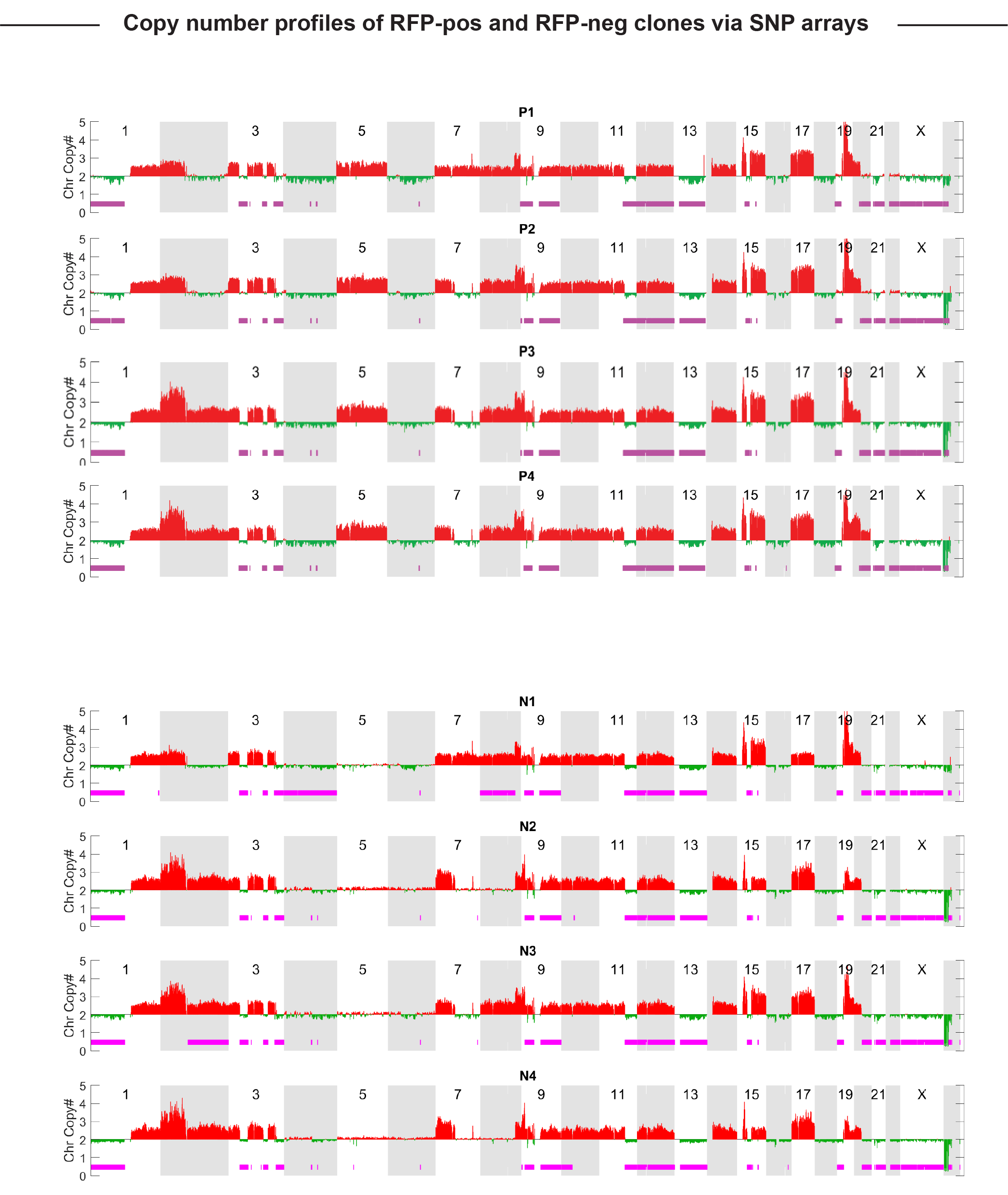
Copy number profiles of RFP-pos and RFP-neg clones via SNP array.

**Figure S4.**
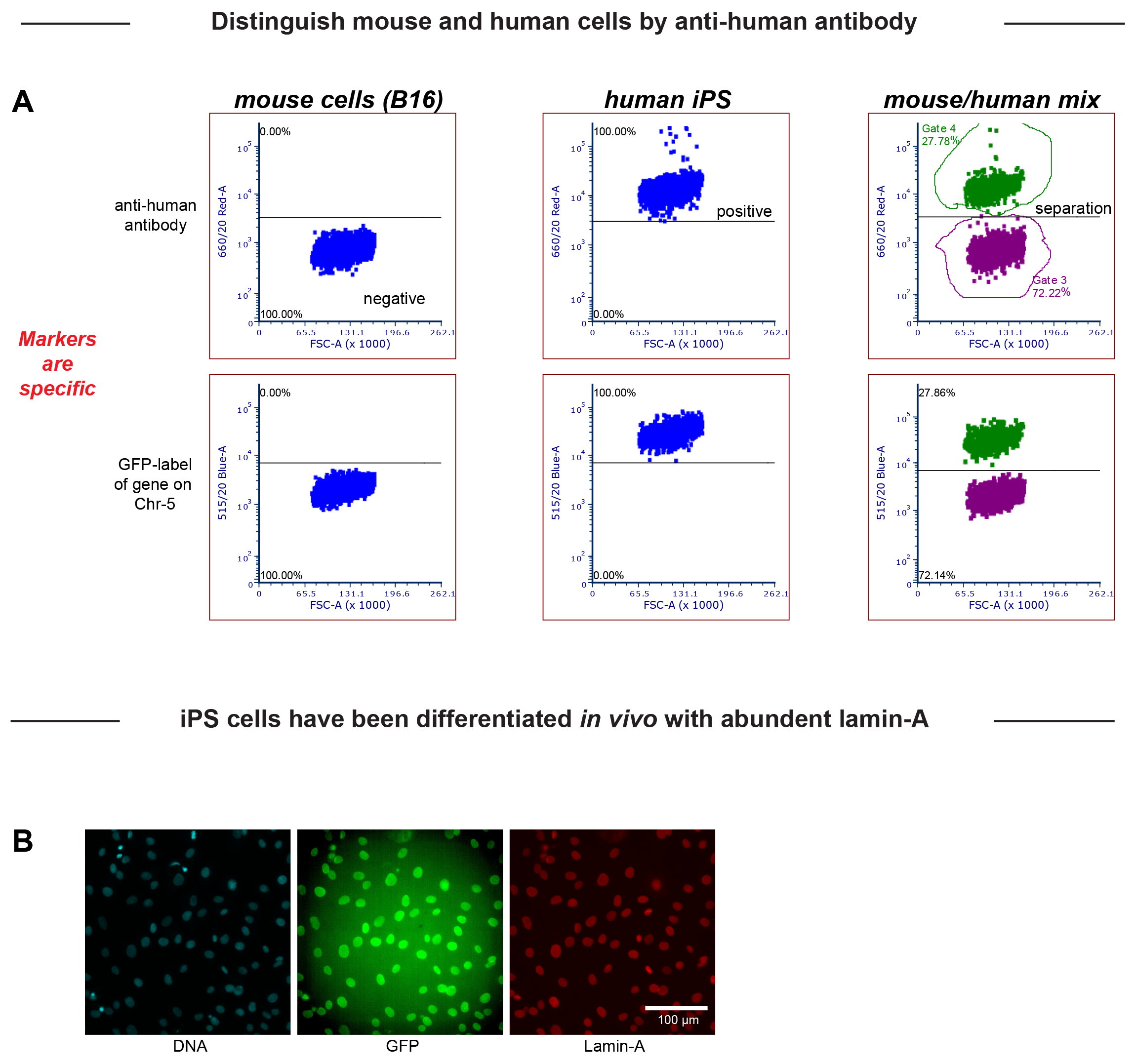
(A) Anti-human antibody is specific to human cells and can be used to distinguish mouse/human cell mixture. (B) iPS cells from teratomas have been differentiated already as high lamin-A is visualized. These iPS cells are proliferating robustly, and their nuclei are larger than mouse cells with higher aspect ratio.

